# Analysis of Plasma Extracellular Vesicles in Normal-Weight and Overweight Type 2 Diabetes Mellitus Using Multimodal SERS and RNA-Seq

**DOI:** 10.64898/2026.03.14.711704

**Authors:** Ugur Parlatan, Aayan Nilesh Patel, Hulya Torun, Asma Humayra Karim, Mehmet Ozgun Ozen, Latha Palaniappan, Utkan Demirci

## Abstract

**Aims:** To characterize subtype-associated heterogeneity in type 2 diabetes mellitus (T2DM), particularly normal-weight diabetes, using extracellular vesicle (EV)-associated molecular features in a clinically stratified cohort.

**Methods:** EVs were isolated from plasma using ExoTIC and validated by transmission electron microscopy, nanoparticle tracking analysis, flow cytometry, and Western blotting. EVs from Asian normal-weight (A-NWD), Asian overweight (A-OWD), Non-Hispanic White normal-weight (W-NWD), and Non-Hispanic White overweight (W-OWD) T2DM patients were analyzed by multimodal surface-enhanced Raman spectroscopy (SERS; n=65) and EV-RNA sequencing (n=39).

**Results:** SERS identified subgroup-associated spectral fingerprints that distinguished the four BMI– and race/ethnicity-defined groups in this cohort. EV-RNA sequencing revealed differential microRNA expression across subgroups, with higher miR-208a and miR-132 in A-OWD and higher miR-484 in A-NWD. Unsupervised analyses also showed partially overlapping EV-associated molecular features between A-NWD and W-OWD, suggesting that BMI-based subgrouping alone may not fully capture shared metabolic states.

**Conclusions:** Multimodal EV profiling identified subgroup-associated spectral and miRNA features in clinically stratified T2DM and provides a framework for studying diabetes heterogeneity, including molecular patterns associated with normal-weight diabetes.

## Introduction

Extracellular vesicles (EVs) are emerging as mediators of metabolic communication and potential signatures for disease heterogeneity in T2DM, which affects 589 million people worldwide^1^. Despite T2DM’s clinical heterogeneity^2^, current classification systems rely primarily on body mass index (BMI), overlooking the molecularly distinct normal-weight diabetes (NWD) phenotype that comprises up to 20% of T2DM patients and paradoxically carries higher mortality risk than overweight/obese T2MD^3^. This phenotype is particularly prevalent in Asian populations who develop T2DM at lower BMI thresholds than other ethnic groups, yet the EV-based molecular signatures underlying these race– and BMI-specific differences remain uncharacterized^4^. Because BMI thresholds may not map equivalently across ancestries, we interpret BMI-stratified comparisons in the context of ancestry-linked differences in body composition and diabetes risk.

EVs facilitate inter-organ communication among insulin-responsive tissues, carrying bioactive cargo, including miRNAs, proteins, and lipids^5,6^, that reflects their cells of origin and modulates recipient cell metabolism^7,8^. In T2DM, circulating EVs from adipose^9^, hepatic^10^, and pancreatic tissues^11^ carry specific miRNA signatures (e.g., miR-208a, miR-132, miR-484) that regulate insulin signaling^12,13^, mitochondrial dynamics^14^, and β-cell function^15^. However, current EV characterization approaches typically examine either cargo composition through sequencing or surface biochemistry through spectroscopy^16–18^ or surface biochemistry through spectroscopy^19–21^, missing the fully integrated picture.

Surface-enhanced Raman spectroscopy (SERS) provides label-free spectral fingerprinting of intact EVs, capturing their protein, lipid, and glycosylation profiles, while RNA sequencing reveals their regulatory cargo^22,23^. Combining SERS with RNA-seq enables parallel assessment of EV-associated spectral phenotype and miRNA cargo from the same EV-enriched preparation. This approach provides enhanced molecular and spectral characterization of T2DM heterogeneity by combining surface and cargo analysis and overcomes the limitations of single-modality analyses that cannot establish such relationships. While prior studies have examined EV protein markers across BMI categories or racial/ethnic groups in diabetes separately, multimodal SERS and RNA-seq analysis to simultaneously characterize label-free biochemical fingerprints and molecular cargo across race– and BMI-stratified T2DM cohorts has not been reported. This integrative strategy leverages EVs’ unique advantages and position them as stable biomarkers.

To address these gaps, we hypothesized that plasma EVs carry distinct molecular cargo reflecting the pathophysiological heterogeneity between normal-weight and overweight T2DM across different racial backgrounds. Given cohort composition and the biological motivation to examine normal-weight diabetes in Asian populations, we pre-specified within-ethnicity comparisons, particularly within the Asian cohort, as primary for inferential RNA analyses, while treating cross-group patterns as cohort-level observations outside the primary inferential analysis. Here, we characterized plasma EVs from 65 T2DM patients stratified by BMI and race/ethnicity using multimodal SERS spectroscopy and miRNA sequencing. As demonstrated in **Fig. 1**, following EV validation, SERS analysis revealed spectral regions that discriminate T2DM subtypes based on lipid and protein signatures, while RNA sequencing identified distinct miRNA profiles: miR-208a/miR-132 upregulation in overweight patients and miR-484 elevation in normal-weight patients, nominating insulin signaling and mitochondrial biology as interpretive themes. Asian normal-weight and White overweight groups showed convergent EV molecular signatures despite different BMIs, suggesting that population-specific BMI thresholds may mask shared EV-associated molecular patterns in this cohort. This multimodal EV profiling strategy enables integrated interpretation of EV-associated spectral and miRNA features across clinically stratified T2DM subgroups.

**Figure 1:**
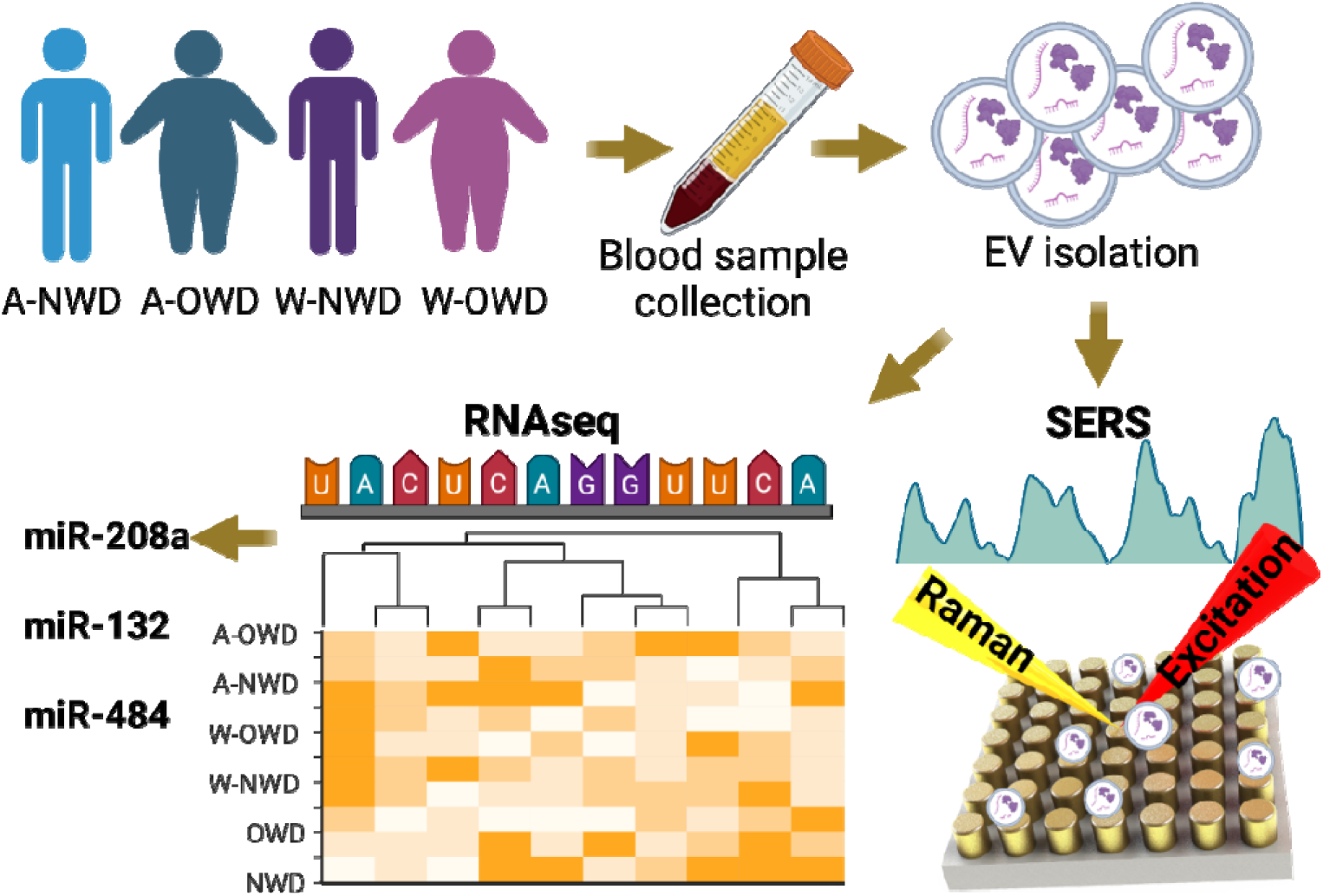
Integrated SERS and RNA-seq workflow for EV profiling in T2DM subtypes. Blood samples from 65 T2DM patients stratified by BMI (normal weight: ≤25 kg/m^2^; overweight: >25 kg/m^2^) and race/ethnicity (Asian [A] or White [W]) were processed to isolate plasma EVs using the ExoTIC method. Purified EVs underwent dual analysis: SERS for label-free spectral fingerprinting (bottom right: Raman spectrum and gold nanostructured substrate) and RNA-seq for miRNA cargo profiling (bottom left: heatmap showing differential expression). Key miRNAs identified include miR-208a and miR-132 (upregulated in A-OWD) and miR-484 (upregulated in A-NWD).

## Results

We first characterized plasma-derived extracellular vesicles to confirm EV identity following MISEV2023 guidelines^24^, assessing size distribution, morphology, and protein markers (**Fig. 2**). To establish EV identity and sample purity via tetraspanin markers already reported^24–26^, we applied bead-captured flow cytometry and Western blotting. Flow cytometry showed high CD63 positivity (97.8% and 95.9%; **Fig. 2a**) and HSPA8 labeling (69.3%), with low calnexin signal (9.3%; **Fig. 2b**), indicating minimal endoplasmic reticulum contamination by cellular debris. To visualize characteristic morphology and size of the EVs, we used TEM and cryo-EM (**Fig. 2c**), which revealed vesicles with cup-shaped/bilayered appearance and diameters within the expected 50–220 nm range. Western blots detected CD63, TSG101, and Flotillin-1, while calnexin was absent (**Supplementary Fig. 1**). To confirm size distribution, we performed nanoparticle tracking analysis (NTA), which showed mode and mean diameters of 97.3 ± 1.9 nm and 126.2 ± 4.2 nm (NWD), and 91.7 ± 1.6 nm and 119.0 ± 4.9 nm (OWD) (**Fig. 2d**). Together, these orthogonal measurements support enrichment for small EVs in the analyzed preparations, while not excluding co-isolated plasma nanoparticles that can overlap in size and composition.

**Figure 2:**
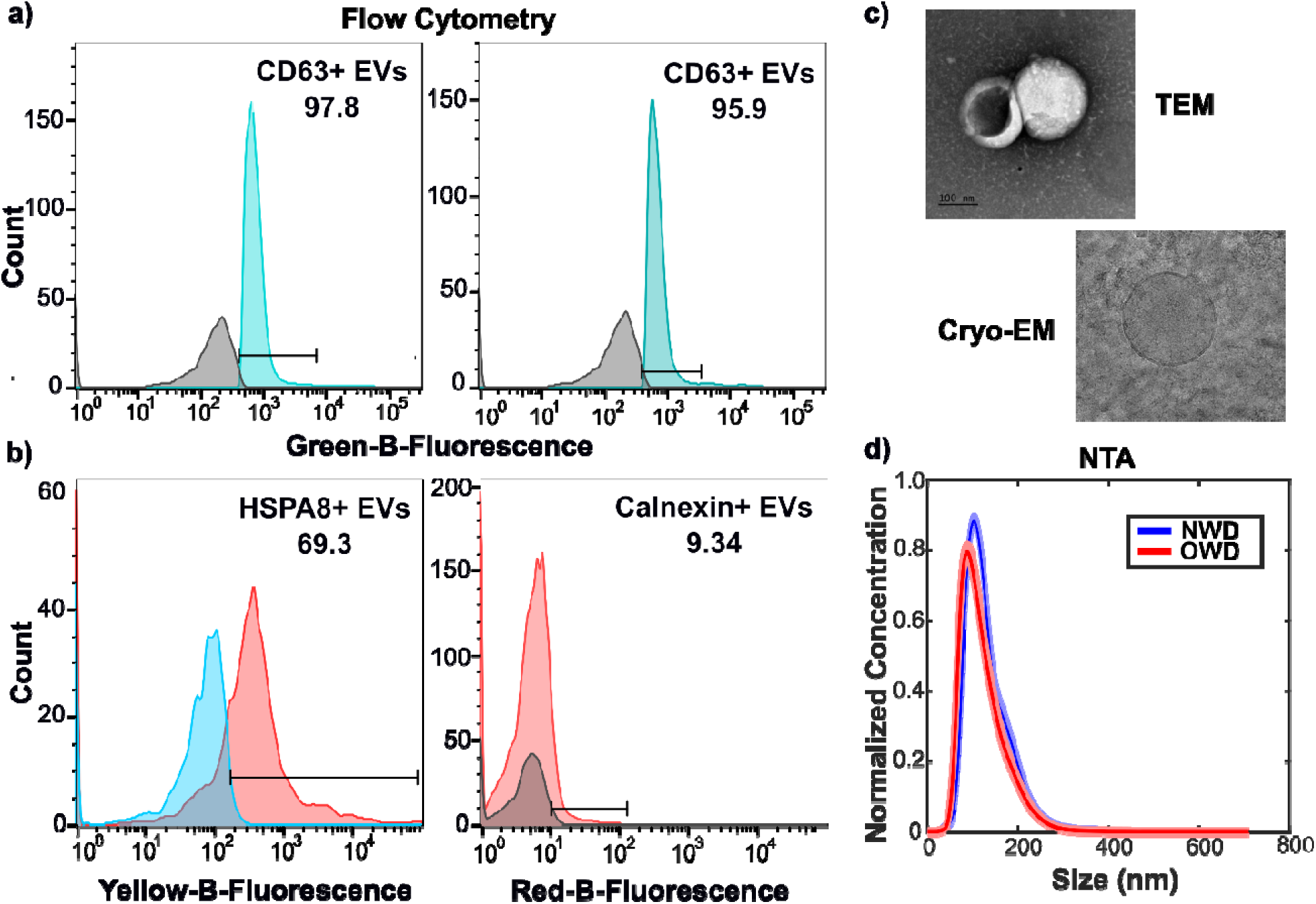
Comprehensive characterization of purified extracellular vesicles (EVs). (a) Representative flow cytometry histograms of EVs immunolabeled with anti-CD63 (turquoise) versus isotype control (gray). Black bars denote the gating region, and percentages indicate the fraction of CD63⁺ particles (97.8% and 95.9%). (b) Flow cytometric detection of HSPA8 (yellow curve; 69.3% positivity) and calnexin (red curve; 9.34% positivity) on EV preparations, demonstrating enrichment of the cytosolic EV marker HSPA8 and minimal contamination by the endoplasmic reticulum protein calnexin. (c) Morphological analysis by electron microscopy. Top: transmission electron micrograph (TEM) showing typical “cup-shaped” EV morphology (scale bar, 100 nm). Bottom: cryo-transmission EM (Cryo-EM) image showing intact lipid bilayer vesicles. (d) Nanoparticle tracking analysis (NTA) size distributions from two independent isolations (blue and red curves), showing mode diameters of 97.3 ± 1.9 nm (NWD) and 91.7 ± 1.6 nm (OWD), consistent with small EV specifications.

Next, we performed SERS analyses on spectral composition of intact EVs derived from four T2DM subgroups, defined by BMI status (normal weight with BMI ≤ 25 kg/m^2^ vs. overweight with BMI > 25 kg/m^2^) and race (Asian vs. Non-Hispanic White). To capture overall spectral differences, we overlaid the average Raman spectra for all groups (**Fig. 3a**), which revealed broad intensity variations across the 800–1600 cm^−1^ range. Because plasma contains abundant non-EV nanoparticles (including lipoproteins^27,28^) with partially overlapping size and biochemical signatures, we interpret SERS features as reflecting the isolated nanoparticle fraction enriched for EV markers rather than asserting EV exclusivity for any single lipid-associated band. Accordingly, band assignments are presented as tentative biochemical interpretations of Raman-active components in the EV-enriched isolates. To identify group-specific spectral shifts, we generated difference spectra (**Fig. 3b**), showing distinct features when both BMI and race were considered. Notably, a signal convergence between the A-NWD and W-OWD groups suggested partially shared EV-associated spectral patterns despite different BMI thresholds across racial backgrounds. This apparent convergence was identified from unsupervised spectral summaries and is presented as a cohort-level pattern rather than evidence of quantitative equivalence between groups. Further, we assessed the statistical significance of the intensity differences between A-NWD and A-OWD using the Mann–Whitney U test and identified wavenumbers meeting the predefined multiple-testing criterion (see Methods) on the difference spectra (**Supplementary Fig. 2**).

**Figure 3:**
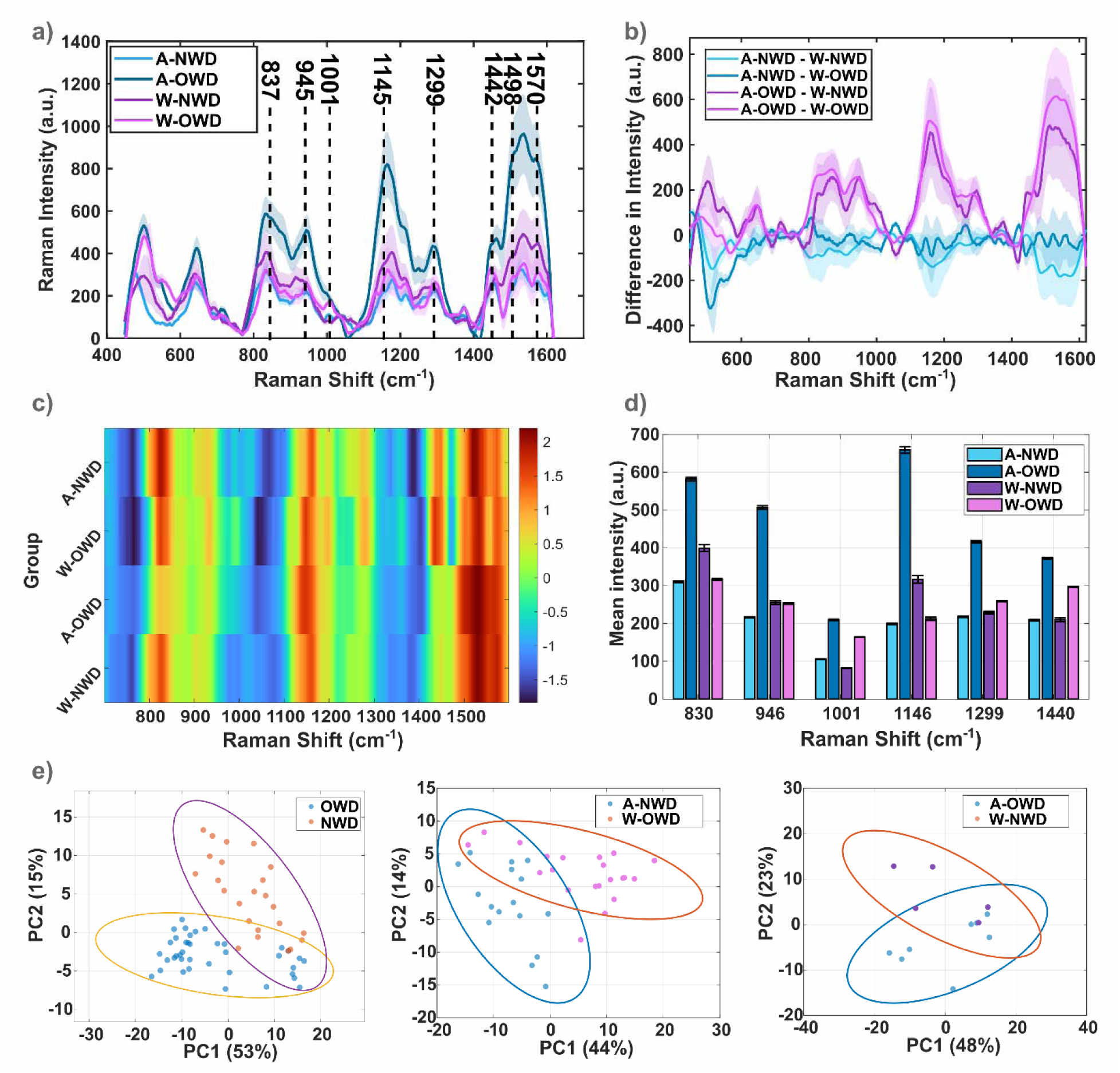
Comparative analysis of SERS data from four T2DM subgroups—Asian Normal Weight Diabetes (A-NWD), Asian Overweight Diabetes (A-OWD), White Normal Weight Diabetes (W-NWD), and White Overweight Diabetes (W-OWD). **(a)** Overlaid average Raman spectra highlight broad differences across the 400–1600 cm^−1^ range. **(b)** Difference spectra (e.g., A-NWD-W-NWD) reveal group-specific shifts. **(c)** A heatmap of average intensities underscores distinct spectral patterns among the four subgroups. **(d)** Bar plot of mean intensity (±SE) at biologically important bands (830, 946, 1001, 1146, 1299, 1440 cm^−1^). **(e)** Patient-wise PCA (PC1–PC2) on normalized spectra within 700–1600 cm^−1^ with 95% covariance ellipses. Left: overall BMI (OWD vs NWD). Middle: Asian NWD vs White OWD. Right: Asian OWD vs White NWD. Axis labels show percent variance explained. Colors are consistent across panels.

To further investigate the molecular bond vibrations that best differentiate these groups, we generated a heatmap of average Raman intensities (**Fig. 3c**). This graph indicates that the three spectral regions exhibit distinct patterns among the different groupings. These region-level differences are used to summarize group-associated spectral variation and do not imply unique molecular assignment of any single band in a complex plasma-derived isolate. The first region, between 785 and 985Lcm^−1^, differentiates BMI scales (NWD vs. OWD) across all races, whereas the other two regions (1130–1346 and 1420–1610Lcm^−1^) are important for distinguishing Asian NWD and Non-Hispanic White OWD groups from the others.

In **Fig. 3d**, we identified statistically significant wavenumbers by performing Mann–Whitney U tests and selected the top 10 significant wavenumbers to construct a bar graph that illustrates the behavior of these peaks among the four groups. Additional statistical comparisons of key spectral markers between NWD and OWD groups, including boxplot distributions (797, 997, 1058, 1256, 1263 cm^−1^), are shown in **Supplementary Fig. 4**. The absolute bond vibrations and tentative biological assignments for these wavenumbers are provided in **Table 1**. Principal component analysis (PCA) score plots (**Fig. 3e**) illustrate clear clustering trends based on both BMI category and racial background. The left panel shows distinct separation between NWD and OWD groups, consistent with BMI-associated spectral variance in EV-enriched plasma nanoparticle isolates. The middle and right panels reveal additional race-specific clustering, where Asian and White subgroups form partially distinct clusters within the same BMI class. Together, these results suggest that both metabolic status (BMI) and racial background contribute to divergence in EV-associated Raman spectral signatures in this cohort.

**Table 1:** Tentative band assignments of the significant peaks.

| Raman shift ($\text{cm}^{-1}$) | Assignment |
| --- | --- |
| **830** | Tyrosine ring breathing (protein)<sup>32</sup> |
| **946** | CCH skeletal stretch or rocking mode (proline, collagen-like)<sup>33</sup> |
| **1001** | Phenylalanine ring breathing (protein)<sup>34</sup> |
| **1146, 1299** | $\text{CH}_2$ twisting/rocking (lipids, membrane components)<sup>35</sup> |
| **1440** | $\text{CH}_2$ bending (lipids, fatty acyl chains)<sup>35</sup> |
| **1498, 1536** | $\text{C}=\text{C}$ stretching (Amide II, some protein modes)<sup>36,37</sup> |
| **1570, 1601** | Aromatic $\text{C}=\text{C}$ stretches (tyrosine, tryptophan)<sup>38,39</sup> |

While SERS revealed distinct surface biochemical fingerprints across T2DM subtypes, these spectral differences could arise from altered EV cargo, biogenesis pathways, or tissue sources. To identify the functional regulatory molecules underlying these biochemical patterns, we performed RNA sequencing on EVs from 39 samples that passed quality control criteria (See **Supplementary Table 1** for patient statistics). We performed differential expression analyses across all subgroup contrasts; however, because statistical power and subgroup balance were strongest within the Asian cohort, we focus statistical interpretation and mechanistic discussion on the Asian NWD versus Asian OWD comparison, while treating other contrasts as exploratory.

In particular, the small W-NWD subgroup size limits stable inference for White within-ethnicity contrasts in the present dataset. **Fig. 4a** shows a heatmap of the top 500 differentially expressed miRNAs across all patient samples, grouped by race/ethnicity and T2DM subtype. Unsupervised clustering revealed that A-OWD samples (green cluster) exhibited a distinct miRNA expression profile compared to NWD groups, with A-OWD samples forming a separate branch and showing the highest expression of multiple miRNAs. Notably, the heatmap revealed similar miRNA expression patterns between A-NWD and W-OWD groups in unsupervised clustering, consistent with the overlapping spectral features in specific lipid-associated regions observed in **Fig. 3c** (1270-1305 cm^−1^ and 1425-1450 cm^−1^).

**Figure 4:**
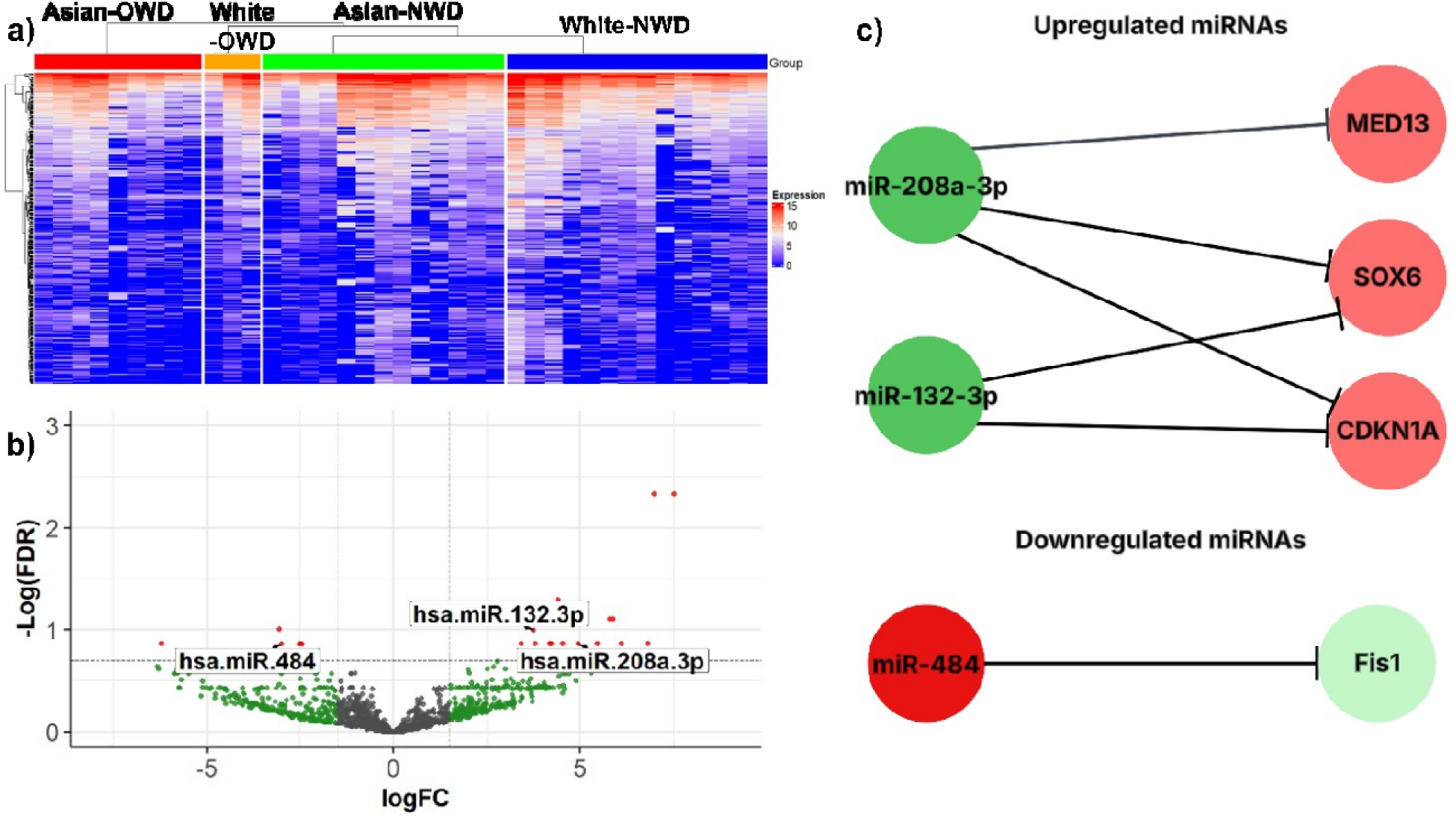
Differential Expression Analysis of Peripheral EV miRNA data. **(a)** Heatmap showing the relative expression of the top 500 differentially expressed miRNAs across all 39 patient samples included in the final cohort, grouped by self-reported race/ethnicity and T2DM subtype. **(b)** A volcano plot showing miRNAs upregulated and downregulated in Asian NWD (reference) and Asian OWD samples. **(c)** Differentially expressed miRNAs between Asian NWD and OWD samples related to T2DM and β-cell pathways and their strongly validated target genes identified through miRTargetLink 2.0. Simplified interaction network shown; full network with all validated targets is provided in **Supplementary Figure 5.**

Stratification by race/ethnicity revealed the most pronounced miRNA dysregulation within the Asian cohort. Therefore, we performed focused differential expression analysis comparing A-NWD and A-OWD to identify population-specific molecular signatures and prioritized biological themes. **Fig. 4b** displays the volcano plot for this comparison, where among 25 candidate differentially expressed miRNAs meeting the exploratory threshold (FDR ≤ 0.2, |logFC| > 1.5), 19 were upregulated and 6 were downregulated in A-OWD relative to A-NWD. Notably, miR-208a-3p and miR-132-3p were significantly upregulated, whereas miR-484 was markedly downregulated in A-OWD. Given the exploratory FDR threshold (FDR ≤ 0.2) and cohort size, these miRNAs are presented as prioritized subgroup-associated miRNAs within the statistical and cohort constraints of this discovery-stage analysis.

Using miRTargetLink 2.0, we mapped the differentially expressed miRNAs to experimentally validated target genes and summarized over-represented biological themes (**Fig. 4c**). KEGG enrichment of validated target genes highlighted diabetes-relevant pathways, including insulin resistance, insulin signaling, Type II diabetes mellitus, PI3K–Akt signaling, glucagon signaling, and AMPK signaling (**Supplementary Table 3**). Within the validated target network, miR-132-3p and miR-208a-3p shared targets implicated in insulin signaling and cell-cycle regulation (including SOX6 and CDKN1A), whereas miR-484 mapped to mitochondrial dynamics regulation via validated targets such as FIS1 (**Supplementary Fig. 5**; **Supplementary Table 3).** These pathway and network outputs provide a data-driven framework for interpreting the observed EV-miRNA associations in the Discussion section.

## Discussion

This study addresses a gap in diabetes heterogeneity research by showing that plasma EV-associated measurements capture race/ethnicity– and BMI-associated molecular signatures not apparent from clinical parameters. Three key findings emerge: (1) SERS and RNA-seq provide complementary EV-associated signatures that distinguish T2DM subgroups in this cohort, (2) Asian NWD and Non-Hispanic White OWD show partially overlapping EV-associated features, and (3) within-Asian miRNA differences nominate insulin signaling and mitochondrial biology themes based on target/pathway mapping.

### EVs as Multimodal Biomarkers for Metabolic Disease Heterogeneity

The EV signatures observed across T2DM subtypes underscore the unique value of EVs as systemic biomarkers. Unlike cell-free analytes such as circulating miRNAs or plasma proteins, EVs integrate molecular information into stable, membrane-protected particles^5,9,10^. The convergence of SERS-based profiling and RNA-seq cargo analysis demonstrates that EVs carry complementary information: lipid and protein composition and internal miRNA content. This dual-layer molecular signature enables more comprehensive disease subtyping than single-modality approaches, positioning plasma EVs as ideal biomarkers for capturing the systemic, multi-organ nature of metabolic disease heterogeneity.

### Plasma EV Spectral Signatures Reveal Population-Specific Differences

The SERS analysis demonstrated that molecular fingerprints at sites associated with fatty acids, glucose, and lipoproteins exhibit distinct race/ethnicity-specific differences as given in **Table 1**. Because several Raman bands in these regions can arise from lipid-rich nanoparticles co-isolated from plasma, we interpret these differences as EV-associated spectral signatures of the analyzed isolates rather than EV-exclusive molecular readouts. The identification of three critical spectral regions (785-985 cm^−1^, 1130-1346 cm^−1^, and 1420-1610 cm^−1^) that differentiate between BMI scales and racial groups suggests that the underlying processes in T2DM are more nuanced than previously understood. Particularly noteworthy is that a lipid-sensitive spectral ratio derived from the 1270 and 1300 cm^−1^ bands (I_1270_ / I_1300_; **Supplementary Fig. 3**)^29^ shows a descriptive inverse trend with group-averaged HbA1c. This finding aligns with previous research indicating that Asian populations may develop T2DM at lower BMI thresholds and exhibit different metabolic profiles compared to other ethnic groups^30,31^.

Similar spectral profiles were observed in Asian NWD and Non-Hispanic White OWD groups (**Fig. 3c**) especially in the bands related to lipids in the regions 1270-1305 cm^−1^ and 1425-1450 cm^−1^ (lipoproteins/fatty acids/triglycerides)^40^, suggesting that comparable molecular alterations may arise despite differences in BMI. This convergence may reflect shared obesity-like EV biochemical signatures in some normal-weight T2DM phenotypes in this cohort.

### Differential EV miRNAs Reflect Distinct Biological Themes Across T2DM Subtypes

RNA sequencing analysis revealed differences in miRNA expression patterns across Asian NWD and OWD patients, providing context for pathway-level interpretation of these T2DM subtypes (**Fig. 4**; **Supplementary Tables 2** and **3**). The upregulation of miR-208a and miR-132 in Asian OWD is consistent with insulin resistance and β-cell dysfunction mechanisms. Both miR-208a and miR-132 converge on SOX6^41,42^ and CDKN1A^43^, which have been implicated in β-cell and cell-cycle biology model systems. miR-208a, which has been shown to regulate systemic energy homeostasis through MED13 in model systems^44^, was significantly upregulated in Asian OWD patients and is discussed here in the context of insulin resistance-related biology. Similarly, miR-132 enhances glucose-stimulated insulin secretion (GSIS)^45^ but may lead to β-cell dysfunction under chronic overexpression conditions^46^. A literature-grounded extended interaction network including additional validated targets (SIRT1, FOXO1, SLC2A1) is provided in **Supplementary Fig. 5,** with full experimental validation details and literature sources in **Supplementary Table 3**.

Because pathway enrichment and validated target mapping are derived from curated interaction resources rather than direct measurements of pathway activity in patient tissues, we describe the miRNA findings in terms of associations with curated targets and enriched pathways. Directionality and physiological interpretations are presented as literature-based context rather than direct inferences from the patient cohort data.

**Table 2:** Functional Characterization of Differentially Expressed miRNAs in Asian T2DM Subtypes.

| miRNA | Expression Pattern | Primary Targets | Biological Process | Pathway Involvement |
| --- | --- | --- | --- | --- |
| miR-208a | ↓ in Asian<br>OWD | MED13<sup>44</sup>,<br>SOX6<sup>42</sup>,<br>CDKN1A<sup>43</sup> | Systemic energy homeostasis, insulin resistance | Cardiac metabolism, β-cell compensation |
| miR-132 | ↓ in Asian<br>OWD | SOX6<sup>41</sup>,<br>CDKN1A<sup>43</sup> | Glucose-stimulated insulin secretion, β-cell proliferation | Insulin signaling, β-cell adaptation |
| miR-484 | ↓ in Asian<br>NWD | Fis1<sup>47</sup> | Mitochondrial fission regulation, ATP production | Mitochondrial dynamics, GSIS |

Conversely, the upregulation of miR-484 in Asian NWD patients and its negative association with obesity suggests a distinct molecular pattern within the Asian NWD subgroup. miR-484 has been reported to regulate mitochondrial network properties via targets such as Fis1 in model systems^47^, and excessive mitochondrial fusion may limit β-cell ATP production critical for GSIS^48^. This finding nominates miR-484 as a potential biomarker for NWD-specific β-cell dysfunction, distinguishing it from the insulin resistance-driven pathology observed in OWD.

These findings are consistent with previous research demonstrating that exosomal miRNAs serve as important mediators of intercellular communication in diabetes^9,12,18^. The differential expression of these miRNAs between NWD and OWD subtypes supports molecular heterogeneity within T2DM beyond BMI-defined categories.

Together, these results define cohort-resolved EV-associated spectral and miRNA patterns that support biological stratification of T2DM heterogeneity beyond BMI alone. Ultimately, translating these molecular signatures into point-of-care diagnostic platforms could support precision management of T2DM subtypes.

## Conclusion

These findings show that plasma EV-associated profiling can resolve molecular patterning within T2DM that is not apparent from BMI alone, including similarities in SERS and RNA signatures between Asian normal-weight and White overweight patients despite different body compositions. Across modalities, the observed EV-associated spectral and miRNA patterns refine biological stratification of T2DM heterogeneity beyond BMI. The identified EV-associated spectral features and subgroup-linked miRNA patterns are interpreted here as cohort-level molecular descriptors that refine biological stratification beyond BMI alone.

## Methods

### Blood Plasma Sample Collection and Extracellular Vesicle (EV) Isolation

Peripheral blood samples were collected from 65 participants (40 from overweight, 25 from normal weight group) under protocols approved by the Institutional Review Board. Detailed baseline characteristics of the study population, including age, sex, race/ethnicity, BMI, and HbA1c levels, are provided in **Supplementary Table 1**. Blood was processed within two hours of collection. Plasma was separated by centrifugation at 1,500LxLg for 10Lminutes to remove cellular components, followed by a 10,000LxLg spin to eliminate debris. Extracellular vesicles were isolated by ExoTIC using the protocol published before^49^. The isolation method includes a washing step to minimize serum protein contamination. The resulting EV sample was resuspended in phosphate-buffered saline (PBS). All purified EV samples were stored at –80°C prior to analysis. Participants were stratified into four clinical subgroups: Asian normal-weight diabetes (A-NWD), Asian overweight diabetes (A-OWD), Non-Hispanic White normal-weight diabetes (W-NWD), and Non-Hispanic White overweight diabetes (W-OWD).

### EV Characterization

EVs were characterized using a combination of Transmission Electron Microscopy (TEM), Cryo-Electron Microscopy (Cryo-EM), Nanoparticle Tracking Analysis (NTA), bead-captured flow cytometry (BC-FC), and Western blotting (WB).

### Bead-Captured Flow Cytometry

EVs were incubated with Aldehyde/Sulfate Latex Beads, 4% w/v, 4 μm ((#A37304, Invitrogen) for 15 minutes at room temperature with gentle rolling. Complexes were blocked with 2% BSA in PBS for 2 hours, followed by glycine (100 mM in PBS) incubation for 30 minutes to minimize nonspecific binding. After thorough washing with cold PBS, bead-EV complexes were incubated overnight with primary antibodies against Calnexin (#PA1-30197, Invitrogen), CD63 (#556019, BD Pharmingen), and HSPA8 (#NB100-41377, Novus Biologicals). The next day, secondary antibodies (#A28175, #A-10931, #31860, Invitrogen) were applied for 2 hours after washing. Following final washes, samples were analyzed on a Guava easyCyte flow cytometer, with data processed using FlowJo v10.

Negative controls including EV-only, bead-only, and bead-plus-EV samples were used to gate the purest EV populations as recommended by MISEV2023. EV populations negative for Calnexin were gated first, followed by multi-parameter gating for CD81– and HSPA8-positive populations as well as double-positive and double-negative subgroups.

### Transmission Electron Microscopy (TEM)

For TEM imaging, EV samples were diluted 10– to 100-fold to optimal concentrations and subjected to negative staining. A 10 µL aliquot of EV suspension was applied to glow-discharged copper grids (Electron Microscopy Sciences, FCF-300-CU) and allowed to adsorb for 3 minutes. The grids were then sequentially stained with three drops of 1% uranyl acetate, each left for 1 minute, followed by removal of excess stain with filter paper. Residual uranyl acetate was removed by three washes with distilled water, blotting between washes. Grids were air-dried for approximately 10 minutes before imaging. EVs were imaged at 100 kV using a JEOL JEM 1400 TEM equipped with a Gatan UltraScan high-resolution digital camera.

### Cryo-EM

High-resolution EV images were obtained by cryo-electron microscopy. Quantifoil Holey Carbon 1.2/1.3 Cu 200 mesh grids were glow-discharged (PELCO easiGlow™, TED PELLA) for 45 seconds at 15 mA. A 3 µL aliquot of EV suspension was applied to the grids, blotted using a Vitrobot Mark IV (FEI) for 3 seconds at blot force 3, with a 10-second wait time, at 22°C and 100% humidity, then plunge-frozen in liquid ethane. Frozen grids were loaded into a ThermoFisher Glacios™ Cryo-TEM operating at 200 kV. Imaging was performed at 36,000× magnification with a pixel size of 1.15 Å using a K3 direct electron detector (Gatan) and SerialEM software. The total electron dose was limited to 60 e⁻/Å², with a defocus range of –1.5 to –3 μm and an exposure time of 3.995 seconds.

### Nanoparticle Tracking Analysis (NTA)

Size distribution and concentration of EVs were quantified by nanoparticle tracking analysis using a Nanosight NS300 system (Malvern Panalytical) with NTA software. A 532 nm green laser (50 mW) was employed, with camera settings optimized at levels 10–14 and detection threshold set to 5. EV samples were diluted 100– to 1000-fold to achieve 20–100 particles per frame.

### Western Blotting

Western blot analysis was performed on EV lysates according to established protocols. Membranes were probed with primary antibodies against Calnexin (Invitrogen, #PA1-30197), TSG101 (Novus Biologicals, #NB200-112), Flotillin-1 (Abclonal, #A6220). HRP-conjugated secondary antibodies, including Goat anti-Rabbit IgG (H+L) (Invitrogen, #31460) and Goat anti-Mouse IgG (H+L) (Invitrogen, #31430), were used as appropriate. All markers were selected in accordance with MISEV2023 recommendations.

### SERS Substrate Preparation and Sample Deposition

Gold-coated nanostructured substrates were fabricated using a template-assisted method to enhance SERS signal uniformity. Prior to use, substrates were rinsed with ethanol, dried under nitrogen, and UV-cleaned for 10Lminutes. A 2 µL aliquot of EV sample was drop-cast onto each substrate and dried at ambient temperature.

### Raman Spectroscopy System

SERS measurements were performed using a custom-built Raman spectroscopy system. A 785 nm diode laser (CrystaLaser; maximum power 130LmW) was used as an illumination source, with power at the sample carefully set below 10LmW using a half-wave plate and linear polarizer to prevent thermal damage. The excitation beam was directed onto the sample through a dichroic mirror (Thorlabs DMLP805) and a 10x microscope objective (NAL=L0.22). Samples were positioned with a motorized XYZ stage (Standa) for precise alignment.

Raman scattering was collected in a 180-degree backscattering geometry using the same objective. Rayleigh scattering was blocked using the dichroic mirror and a Semrock ultra-steep long-pass filter. The collected signal was coupled into a multi-mode fiber and detected with a StellarNet HyperNova spectrometer equipped with a –60°C-cooled CCD camera (Andor iVac 316) to minimize dark noise.

Multiple spatially distinct points were sampled per EV specimen to address spatial heterogeneity. At each point, a 10-second integration time was used, and three consecutive scans were averaged per measurement.

### Spectral Processing and Statistical Analysis

All spectral processing was conducted using a custom pipeline in MATLAB. Raw spectra were first calibrated using a polystyrene reference. Smoothing was applied with a Savitzky-Golay filter (second-order polynomial, 11-point window), followed by baseline correction via the Asymmetric Least Squares method with standardized parameters. All spectra were vector-normalized (L2 norm), aligned to a common wavenumber grid (500–1800 cm^−1^, 1 cm^−1^ steps), and outlier spectra were excluded using interquartile range filtering on total spectral intensity.

#### Statistical Analysis

Pairwise group comparisons were performed at each wavenumber using the Mann–Whitney U test. Significance was defined as Bonferroni-corrected p-valuesL<L0.01. Significant wavenumbers were interpreted in the context of reported biochemical bond vibrations in complex biological samples.

#### Principal Component Analysis

PCA was performed using MATLAB’s *pca()* function on all preprocessed spectra. The first two principal components (PC1 and PC2), which together captured the dominant variance structure in the dataset, were used for visualization of group clustering.

#### Data Visualization

Normalized intensities of significant peaks were visualized using heatmaps (z-scored per group) and bar plots (key wavenumbers with group-wise means ± standard error), to illustrate biochemical variation across study cohorts.

### RNA Extraction

EV-enriched isolates were obtained from human peripheral blood using the ExoTIC workflow described above, and RNA was subsequently isolated with the miRNeasy Serum/Plasma Kit (QIAGEN). Briefly, EV pellets were disrupted in QIAzol Lysis Reagent, homogenized, and subjected to chloroform-induced phase separation. The RNA-containing aqueous phase was combined with ethanol and transferred to RNeasy Mini spin columns. After sequential washes with Buffers RWT and RPE to eliminate impurities, total RNA—including small RNAs longer than 18 nucleotides—was eluted in RNase-free water. RNA quality and concentration were evaluated with the Agilent 2100 Bioanalyzer using the Small RNA Assay. All extractions were processed in a single batch to minimize variability between runs.

### RNA Sequencing and Data Processing

Small RNA libraries were prepared from the extracted RNA and sequenced on an Illumina platform, producing 75 bp single-end reads. Raw data were processed using the EV Small RNA-seq Analysis 1.9 pipeline (Banana Slug Genomics Center, UCSC). All RNA sequencing was performed in a single batch to avoid batch effects.

Subsequently, reads were analyzed with the nf-core small RNA-seq pipeline (v2.4.0; https://nf-co.re/rnaseq/3.14.0/). Quality control was carried out using FastQC (v0.12.0), while adapter trimming and low-quality read filtering were performed with fastp (v0.23.4). Barcode deduplication utilized UMI-tools (v1.1.4). For miRNA-specific quality assessment and origin tracking, miRTrace (v1.0.1; https://github.com/friedlanderlab/mirtrace) was applied. Reads were aligned with Bowtie1 (v1.2.2) against a custom index based on the miRBase database (https://www.mirbase.org/), followed by post-processing with Samtools (v1.21). Mature and precursor miRNA counts were generated using edgeR (v3.20), and annotation was performed with mirtop (v0.4.28; https://github.com/miRTop/mirtop). Novel and known miRNA discovery was completed with MiRDeep (v0.1.3; https://github.com/rajewsky-lab/mirdeep2). Quality control results were summarized and visualized with MultiQC (v1.25).

### Differential Expression Analysis

Differential expression was conducted in R using the edgeR package (v4.2.2, Bioconductor), based on the established workflow of Robinson et al. (2010). Raw miRNA counts were formatted into a DGEList object with sample metadata included. Genes with very low counts were removed prior to analysis to minimize noise. To adjust for library size and composition, normalization was performed using the trimmed mean of M-values (TMM) method.

Dispersion estimates were modeled with negative binomial generalized linear models (GLMs) to compare expression profiles between participants in the overweight group (BMI > 25) and the normal weight group (BMI <= 25), both across the full dataset and within individual ethnic subgroups (Figure 1). Within-ethnicity analyses were prioritized in cases of unequal ethnic representation between OWD and NWD groups. Hypothesis testing was carried out using quasi-likelihood F-tests within the GLM framework. P-values were corrected for multiple hypothesis testing using the Benjamini–Hochberg procedure to control the false discovery rate (FDR). Differential expression was considered significant at FDR ≤ 0.2 and |logFC| > 1.5. We computed differential expression for all groups, but pre-specified the Asian NWD vs Asian OWD comparison for formal inference because it provided the most balanced subgroup sizes and statistical power; other contrasts are reported as exploratory. Significance was visualized with volcano plots generated by the EnhancedVolcano package (v1.22.0), and overall miRNA expression patterns were shown with heatmaps created using the ComplexHeatmap package (v2.20.0).

### Gene Ontology and Pathway Analysis

Differentially expressed miRNAs were separated into upregulated and downregulated groups relative to the NWD reference group. These miRNAs were analyzed with miRTargetLink 2.0 to identify experimentally validated gene targets, and only miRNAs associated with at least two strong target-gene interactions were retained for downstream analyses. Enrichment analyses of target genes were performed using GeneTrail 3.0 and Kyoto Encyclopedia of Genes and Genomes (KEGG) pathways. Results were further contextualized through a literature review of pathways previously implicated in T2DM and nutritional weight differences.

## Acknowledgements

U.D. acknowledges the support of a seed award from Stanford University Diabetes Research Center. L.P. acknowledges funding from the National Institutes of Health under the project number 1K24HL150476.

## Author contributions

*Conceptualization:* U.P, L.P., and U.D.; *Methodology:* U.P, A.N.P., H.T., M.O.O., A.H.K., L.P. and U.D*; Software*: U.P and A.N.P.; Data Curation, *Validation and Formal analysis:* U.P, A.N.P, A.H.K., M.O.O. and H.T.; *Investigation:* U.P, A.N.P., M.O.O, A.H.K. and H.T.; *Writing-Original Draft*: U.P, A.N.P, H.T., M.O.O., L.P. and U.D; *Writing-Review and Editing*: U.P, A.N.P, H.T., M.O.O., A.H.K., L.P. and U.D*; Supervision, Project administration, Funding acquisition and Resources:* L.P. and U.D.

## Data availability

All data supporting the findings of this study are available within the paper and its Supplementary Information. The custom codes used to generate Figures 2 and 3 and the related data was uploaded on Zenodo with the identifier https://doi.org/10.5281/zenodo.18945379.50

Additional data that support the findings of this study are available from the corresponding author upon request.

## Declaration of generative AI and AI-assisted technologies in the manuscript preparation process

During the preparation of this work the authors used ChatGPT in order to improve the flow of the text. After using this tool, the authors reviewed and edited the content as needed and take full responsibility for the content of the published article.

