## Supplementary Fig for "Analysis of Plasma Extracellular Vesicles in Normal-Weight and Overweight Type 2 Diabetes Mellitus Using Multimodal SERS and RNA-Seq"

1. **Patient statistics – miRNA Statistical Analysis**

**Supplementary Table 1:** Baseline characteristics of Asian and Non-Hispanic White patients from NWD and OWD groups in final RNA-seq cohort (n = 39)

|  | **Normal-Weight Diabetes** | **Overweight Diabetes** |  |
| --- | --- | --- | --- |
| **N** | 16 | 23 |  |
| **Mean Age (SD)** | 61.5 (10.9) | 58.4 (9.2) | p = 0.37 |
| **Male Sex (%)** | 11 (68.8%) | 17 (73.9%) | p = 0.73 |
| **Race/Ethnicity (%)** | | |  |
| **Non-Hispanic White** | 3 (18.8%) | 14 (60.9%) | p = 0.018 |
| **Asian** | 13 (81.3%) | 9 (39.1%) |  |
| **Mean BMI (SD)** | 23.8 (1.3) | 35.4 (6.8) | p < 0.001 |
| **Mean HbA1c (SD)** | 7.8 (0.9) | 7.8 (1.2) | p = 0.87 |

2. **EV Characterization – Western Blots**

To confirm the successful enrichment of EVs from plasma samples across the study cohort (representing diverse BMI characteristics), we performed Western blot analysis on the isolated fractions. As shown in Supplementary Figure S1, the samples showed strong positive expression for the cytosolic ESCRT-related proteins TSG101 and Flotillin-1, which are canonical markers of EVs and small EVs. Importantly, the samples were negative for Calnexin, an endoplasmic reticulum protein not expected to be sorted into EVs. The absence of Calnexin indicates that the isolation protocol minimized detectable intracellular contamination in this assay, supporting the use of EV-enriched isolates for downstream biomarker analyses.
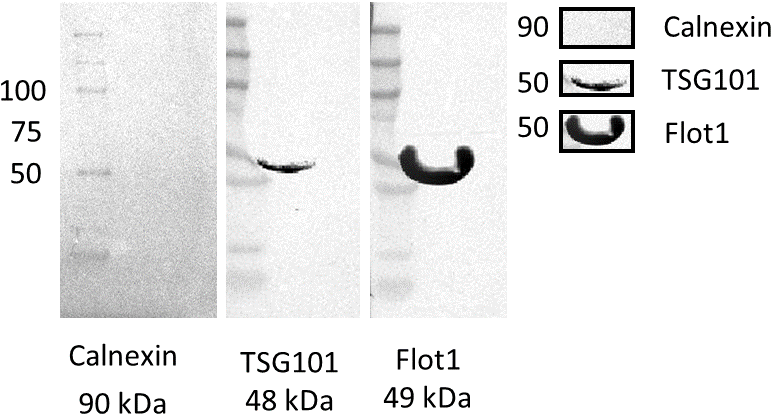

**Supplementary Figure 1: Western blot validation of extracellular vesicle (EV) isolation from human plasma.** The isolated vesicles show enrichment of the classical EV markers TSG101 (48 kDa) and Flotillin-1 (Flot1) (49 kDa). The endoplasmic reticulum marker Calnexin (90 kDa) is absent, supporting enrichment of EV-associated markers and reduced detectable cellular contamination in the analyzed isolates. Molecular weight markers (kDa) are indicated on the left. Band shape variations reflect the nature of the gel matrix and loading conditions but do not affect the qualitative validation of marker presence/absence.

3. **SERS characterization**

To identify specific biochemical alterations associated with obesity in diabetic patients, we calculated the difference spectrum by subtracting the mean signal of the Normal Weight Diabetic (NWD) group from the Overweight Diabetic (OWD) group (OWD - NWD). As shown in **Supplementary Figure 2**, statistically significant spectral differences (FDR < 0.05) were identified across multiple bands in the fingerprint region (800–1600 cm^-1^).

**Lipid Enrichment in OWD**: The OWD group exhibited higher intensities (positive peaks) in regions characteristic of lipids, particularly around 1060–1130 cm^-1^ (C-C stretching) and 1440–1460 cm^-1^ (CH_2_/CH_3_ deformation), suggesting an increased lipid load or specific lipid subclass enrichment in EVs from overweight patients.

**Protein Variations:** Significant negative features were observed in the amide III region (1240–1280 cm^-1^) and the tyrosine doublet region (830–850 cm^-1^), indicating structural or compositional changes in EV proteins associated with the normal weight phenotype compared to the overweight phenotype.

**Tyrosine/Phenylalanine**: The sharp decrease around 1003 cm^-1^ (phenylalanine ring breathing) in the OWD group relative to NWD suggests a relative reduction in phenylalanine content or signal suppression in the overweight cohort.

Together, these biochemical differences support BMI-associated variation in EV-associated Raman signatures measured from EV-enriched plasma isolates, complementing clinical parameters such as HbA1c in this cohort. Band assignments in this section are used as tentative biochemical annotations of Raman-active components in a complex plasma-derived isolate and are not intended as definitive attribution to EV membrane constituents alone.

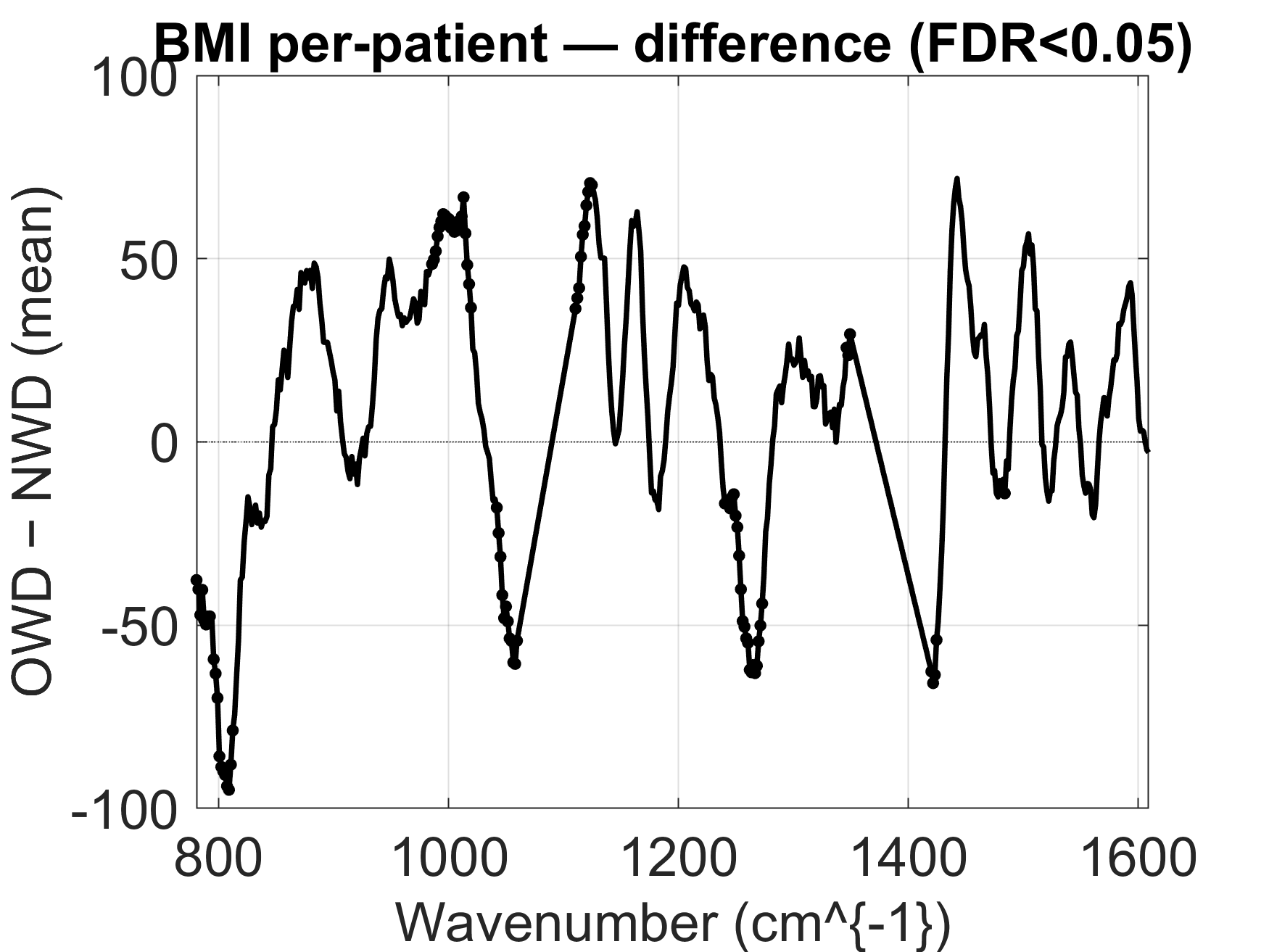

**Supplementary Figure 2**: **Difference spectrum of plasma EVs between Overweight Diabetic (OWD) and Normal Weight Diabetic (NWD) patients**. The plot displays the mean difference spectrum calculated by subtracting the NWD mean spectrum from the OWD mean spectrum (OWD - NWD) in the fingerprint region (800–1600 cm^-1^). Black markers indicate wavenumbers with statistically significant differences (False Discovery Rate, FDR < 0.05). Positive peaks represent spectral features enriched in the OWD group, while negative peaks correspond to features enriched in the NWD group. Major differences are observed in bands commonly annotated to lipid- and protein-associated vibrations in complex biological samples (e.g., ~1440 cm^-1^, ~1060-1100 cm^-1^, 1240-1280 cm^-1^, ~800-850 cm^-1^).

4. **HbA1c-SERS relationship**

Across subgroup averages, I_1270_ / I_1300_ exhibits a qualitative inverse trend with mean HbA1c (**Supplementary Fig. 3**). We interpret this observation conservatively as a descriptive subgroup-level association, consistent with HbA1c-associated differences in lipid-sensitive Raman features in EV-enriched plasma isolates, without implying patient-level prediction or causal mechanistic links. Because this ratio is a composite spectral feature in a complex plasma-derived isolate, it is not interpreted as an EV-exclusive molecular readout. The direction of this subgroup trend is consistent with literature reporting glycemia-associated changes in lipid-related Raman features.
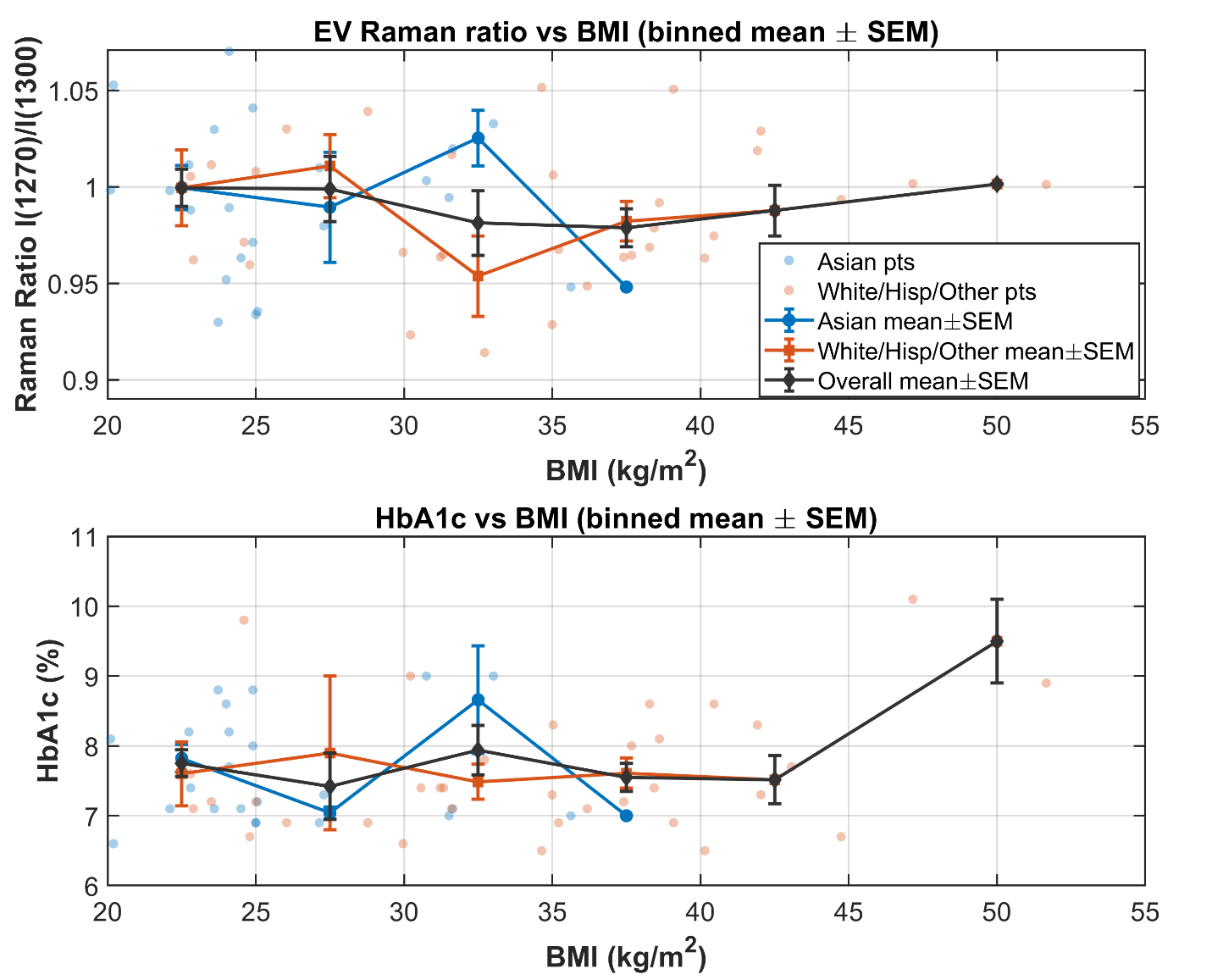

**Supplementary Figure 3:** **Group-level relationship between HbA1c and a lipid-sensitive SERS ratio. Top panel:** EV Raman intensity I1270/I1300 plotted against BMI. **Bottom panel**: HbA1c (%) plotted against BMI. In each panel, faint points indicate individual patients, while connected markers show BMI-binned group means for Asian patients, White/Hispanic/Other patients, and the overall cohort. Error bars represent SEM (standard error of measurement). The 1270 cm^-1^ band is commonly associated with unsaturated lipid-related =C–H in-plane deformation, whereas the 1300 cm^-1^ band is attributed to CH_2_ twisting/bending in lipid acyl chains.

5. **Comparison of average SERS intensities at selected discriminatory wavenumbers**

To pinpoint the specific molecular alterations driving the separation between BMI groups, we performed a statistical comparison of peak intensities at distinct wavenumbers. As illustrated in **Supplementary Figure 4**, the Overweight Diabetic (OWD) group exhibited a highly significant increase in the intensity of the band centered at 1002 cm^-1^ (averaged between 997 and 1005 cm^-1^, p = 7.1x10^-10^), which is assigned to the ring breathing mode of phenylalanine. This elevation is consistent with increased intensity of a phenylalanine-associated Raman feature in EV-enriched isolates from overweight patients.

Conversely, the Normal Weight Diabetic (NWD) group showed significantly higher intensities in the ~797 cm^-1^ band (p = 2.8x10^-8^), assigned to the O-P-O backbone stretching of nucleic acids (DNA/RNA), and the ~1058 cm^-1^ band (p = 2.4x10^-6^), characteristic of C-C stretching in lipids. Additionally, the NWD group displayed stronger signals in the Amide III region at ~1256 cm^-1^ and ~1263 cm^-1^ (p < 0.001), consistent with differences in protein-associated Raman features between the two cohorts. The band at ~1482 cm^-1^ (attributed to guanine/adenine) followed a similar trend (higher in NWD) but was not statistically significant (p = 0.071).

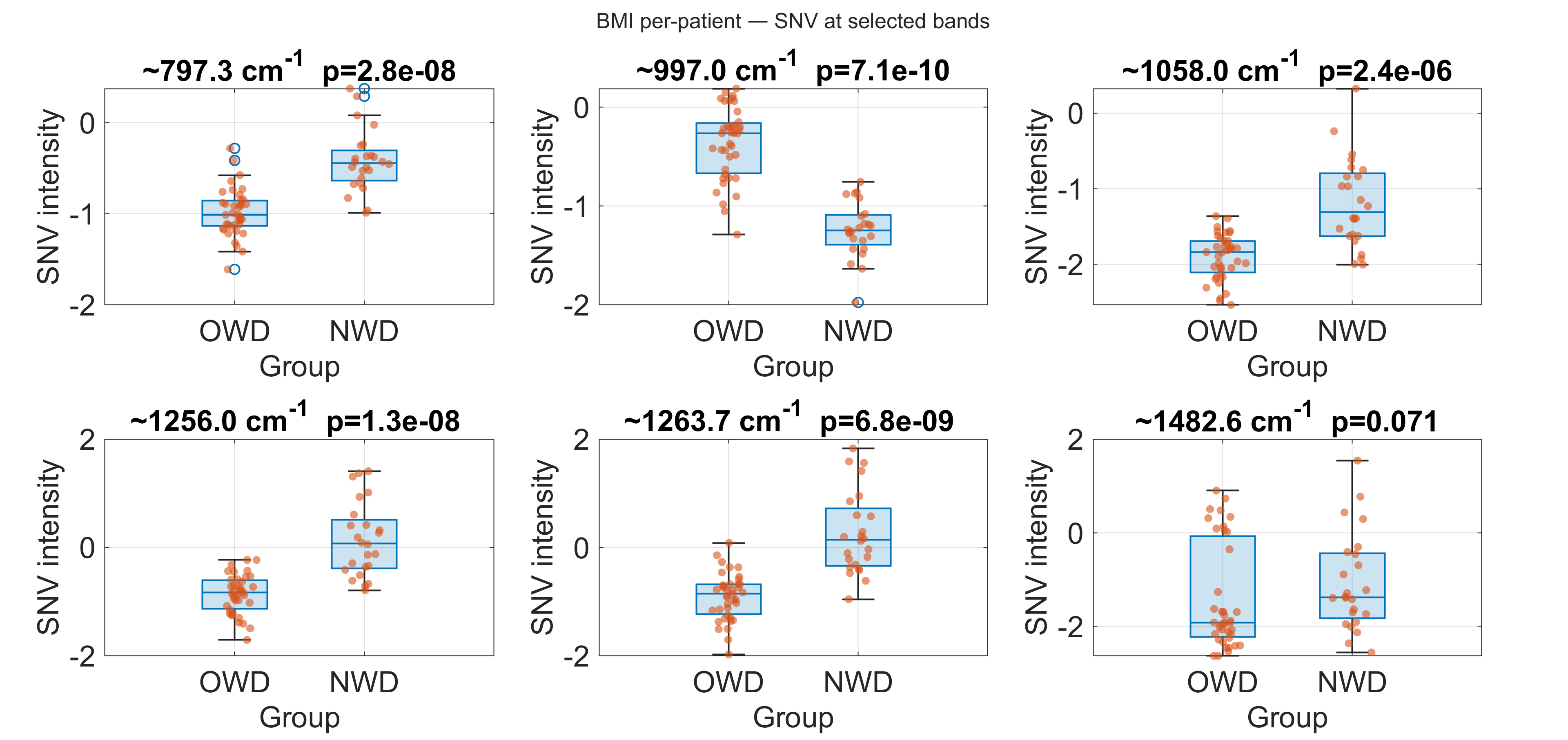

**Supplementary Figure 4: Statistical comparison of key spectral markers between Overweight Diabetic (OWD) and Normal Weight Diabetic (NWD) groups.** Boxplots showing normalized intensities of selected discriminatory wavenumbers in spectra measured from EV-enriched plasma isolates. The OWD group is compared to the NWD group. Significant differences were observed at ~797 cm^-1^ (DNA/RNA backbone), ~997 cm^-1^ (Phenylalanine), ~1058 cm^-1^ (Lipid C-C stretch), and the Amide III region (~1256 cm^-1^ and ~1263 cm^-1^). The band at ~1482 cm^-1^ (DNA/RNA, Guanine/Adenine) showed a trend but did not reach statistical significance (p = 0.071). P-values (Mann-Whitney U test) are displayed above each plot. Relative to NWD, the OWD group shows higher intensity at the phenylalanine-associated ~997 cm^-1^ feature and lower intensity at several other wavenumbers, including ~797 cm^-1^ and the Amide III region (~1256/1263 cm^-1^).

6. **Extended miRNA target and pathway annotation network**

To provide a broader interpretive view of the differentially expressed miRNAs, we constructed an extended interaction network (**Supplementary Figure 5**) mapping miR-132-3p, miR-208a-3p, and miR-484 to their experimentally validated targets involved in glucose homeostasis, insulin signaling, and mitochondrial dynamics.

This network expands upon the summarized pathways presented in the main text by including a broader range of targets such as SIRT1, FOXO1, and SLC2A1 (targets of miR-132-3p), which are critical for insulin sensitivity and metabolic stress response. In curated target annotations, miR-208a-3p includes MED13 among validated targets relevant to systemic energy homeostasis, while miR-484 includes FIS1, a validated target involved in mitochondrial dynamics. These relationships are presented as literature-based context for interpreting the observed EV-miRNA patterns. Full details of the validated miRNA-target interactions, including the experimental methods (e.g., Luciferase reporter assays, Western blot) and literature sources (PMIDs) used to construct this network, are provided in **Supplementary Table 3** below.

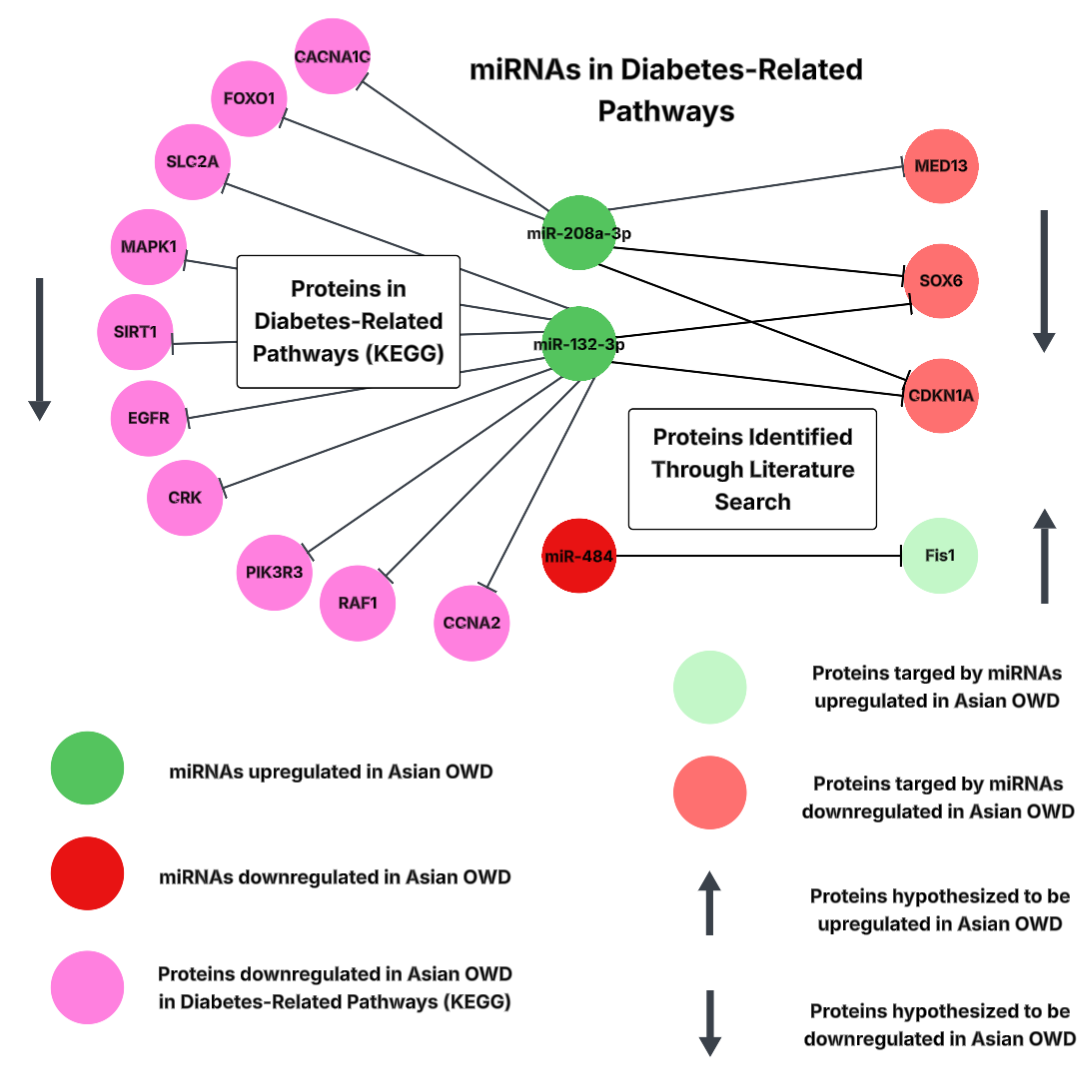

**Supplementary Figure 5: Extended interaction network of differentially expressed miRNAs and their gene targets in diabetes pathways.** The network diagram illustrates the comprehensive regulatory relationships between the key miRNAs identified in this study (miR-132-3p, miR-208a-3p, miR-484) and their validated target genes associated with diabetes, metabolic regulation, and mitochondrial function. (Left) miR-208a-3p (upregulated) targets critical metabolic regulators including MED13 (Mediator of RNA polymerase II transcription subunit 13) and SOX6, which regulate glucose and lipid metabolism, as well as the cell cycle inhibitor CDKN1A (p21). (Center) miR-132-3p (upregulated) exhibits a dense network of targets involved in insulin signaling and cell survival, such as SIRT1, FOXO1, MAPK1, PIK3R3, and the glucose transporter SLC2A1. It also shares the target SOX6 and CDKN1A with miR-208a-3p, suggesting potential cooperative regulation. (Right) miR-484 (downregulated) is linked to its primary target FIS1 (Mitochondrial fission 1 protein), a key regulator of mitochondrial dynamics and apoptosis. Green nodes represent upregulated miRNAs; Red nodes represent downregulated miRNAs; Pink/Red target nodes indicate experimentally validated targets (based on literature) that are hypothesized to be downregulated or upregulated, respectively, in response to miRNA changes. Network connections summarize curated, experimentally validated miRNA–target relationships from external databases/literature and are not direct measurements of pathway activity in patient tissues.

**Supplementary Table 2:** Gene targets and source information for differentially expressed miRNAs between NWD and OWD Asian patient samples

| **miRNA** | **Target / Target pathway** | **Source** | **Experiment(s)** | **Reference - PMID** |
| --- | --- | --- | --- | --- |
| **hsa-miR-1266-5p** | TERT | MIRT053598 | Immunohistochemistry, Luciferase reporter assay, Microarray, qRT-PCR, Western blot | 24481448 |
| **hsa-miR-132-3p** | SIRT1 | MIRT000333 | Luciferase reporter assay, Western blot | 19819989 |
| **hsa-miR-132-3p** | SIRT1 | MIRT000333 | ChIP-seq, Immunoprecipitation, Luciferase reporter assay, Microarray, qRT-PCR, Western blot | 28383763 |
| **hsa-miR-132-3p** | CDKN1A | MIRT000601 | qRT-PCR, Luciferase reporter assay, Western blot | 20190813 |
| **hsa-miR-132-3p** | RB1 | MIRT006503 | Luciferase reporter assay, Western blot, Reporter assay;Western blot;qRT-PCR | 21329664 |
| **hsa-miR-132-3p** | ARHGAP32 | MIRT003926 | Luciferase reporter assay, Western blot | 18577589 |
| **hsa-miR-132-3p** | ARHGAP32 | MIRT003926 | Luciferase reporter assay | 19850129 |
| **hsa-miR-132-3p** | TLN2 | MIRT007253 | Luciferase reporter assay | 22310291 |
| **hsa-miR-132-3p** | RASA1 | MIRT007010 | Immunoblot, Western blot | 21868695 |
| **hsa-miR-132-3p** | HBEGF | MIRT006659 | Luciferase reporter assay, Microarray, qRT-PCR, Western blot | 21853268 |
| **hsa-miR-132-3p** | HBEGF | MIRT006659 | Luciferase reporter assay | 22310291 |
| **hsa-miR-132-3p** | CRK | MIRT007166 | Luciferase reporter assay | 23184980 |
| **hsa-miR-132-3p** | TJAP1 | MIRT007167 | Luciferase reporter assay | 23184980 |
| **hsa-miR-132-3p** | TLN2 | MIRT007253 | Luciferase reporter assay | 22330136 |
| **hsa-miR-132-3p** | SPRED1 | MIRT021849 | Luciferase reporter assay, Western blot | 25945589 |
| **hsa-miR-132-3p** | MUC13 | MIRT021860 | Immunohistochemistry, Luciferase reporter assay, qRT-PCR, Western blot | 28339011 |
| **hsa-miR-132-3p** | CCNA2 | MIRT021865 | Western blot;qRT-PCR | 21329664 |
| **hsa-miR-132-3p** | CCNB1 | MIRT021866 | Western blot;qRT-PCR | 21329664 |
| **hsa-miR-132-3p** | BDNF | MIRT054502 | qRT-PCR, Western blotting | 23704927 |
| **hsa-miR-132-3p** | FOXO1 | MIRT045840 | Luciferase reporter assay, qRT-PCR, Western blot | 26298723 |
| **hsa-miR-132-3p** | FOXO1 | MIRT045840 | Luciferase reporter assay, Western blot | 27556448 |
| **hsa-miR-132-3p** | SOX5 | MIRT054894 | Luciferase reporter assay, Western blot | 25305446 |
| **hsa-miR-132-3p** | JPT1 | MIRT054835 | Luciferase reporter assay, qRT-PCR, Microarray, Western blot | 25538559 |
| **hsa-miR-132-3p** | KLHL11 | MIRT054836 | Luciferase reporter assay, qRT-PCR, Microarray, Western blot | 25538559 |
| **hsa-miR-132-3p** | MAPK1 | MIRT054837 | Luciferase reporter assay, qRT-PCR, Microarray, Western blot | 25538559 |
| **hsa-miR-132-3p** | IRAK4 | MIRT438882 | Luciferase reporter assay | 23264652 |
| **hsa-miR-132-3p** | SOX4 | MIRT732559 | Immunoprecipitation, Luciferase reporter assay, qRT-PCR, Western blot | 26377202 |
| **hsa-miR-132-3p** | EGFR | MIRT731713 | Immunohistochemistry, Luciferase reporter assay, qRT-PCR, Western blot | 26452132 |
| **hsa-miR-132-3p** | RAF1 | MIRT731714 | Immunohistochemistry, Luciferase reporter assay, qRT-PCR, Western blot | 26452132 |
| **hsa-miR-132-3p** | SPRY1 | MIRT733351 | Luciferase reporter assay, Western blot | 25945589 |
| **hsa-miR-132-3p** | YY1AP1 | MIRT733928 | Luciferase reporter assay, qRT-PCR, Western blot | 26096363 |
| **hsa-miR-132-3p** | MMP9 | MIRT733764 | Luciferase reporter assay | 26319558 |
| **hsa-miR-132-3p** | AGO2 | MIRT734062 | Immunoblot, Luciferase reporter assay, qRT-PCR | 26475020 |
| **hsa-miR-132-3p** | PIK3R3 | MIRT734832 | Luciferase reporter assay, qRT-PCR, Western blot | 26252738 |
| **hsa-miR-132-3p** | SOX6 | MIRT734812 | Luciferase reporter assay, Western blot | 27556448 |
| **hsa-miR-132-3p** | GDF5 | MIRT734813 | Luciferase reporter assay, Western blot | 27556448 |
| **hsa-miR-132-3p** | SMAD2 | MIRT735158 | GFP reporter assay, qRT-PCR, Western blot | 25988335 |
| **hsa-miR-132-3p** | SLC2A1 | MIRT735435 | Luciferase reporter assay, qRT-PCR, Western blot | 27398313 |
| **hsa-miR-208a-3p** | MED13 | MIRT003912 | Luciferase reporter assay, qRT-PCR, Western blot | 17379774 |
| **hsa-miR-208a-3p** | CDKN1A | MIRT000599 | qRT-PCR, Luciferase reporter assay, Western blot | 20190813 |
| **hsa-miR-208a-3p** | CDKN1A | MIRT000599 | Luciferase reporter assay, Western blot | 26754670 |
| **hsa-miR-208a-3p** | ETS1 | MIRT005541 | Luciferase reporter assay, Microarray, qRT-PCR, Western blot | 20576608 |
| **hsa-miR-208a-3p** | SOX6 | MIRT438545 | Luciferase reporter assay, Western blot | 25023649 |
| **hsa-miR-208a-3p** | CACNA1C | MIRT732875 | Luciferase reporter assay | 27545043 |
| **hsa-miR-208a-3p** | CACNB2 | MIRT732877 | Luciferase reporter assay | 27545043 |
| **hsa-miR-208a-3p** | PDCD4 | MIRT733324 | Luciferase reporter assay, qRT-PCR, Western blot | 27634902 |
| **hsa-miR-208a-3p** | Qk | MIRT734241 | Luciferase reporter assay | 28283792 |
| **hsa-miR-208a-3p** | QKI | MIRT734998 | Luciferase reporter assay | 28283792 |
| **hsa-miR-4496** | ABCG2 | MIRT733729 | Luciferase reporter assay, Western blot | 28008607 |
| **hsa-miR-484** | FIS1 | MIRT006714 | ChIP-seq, Immunoblot, LacZ reporter assay, qRT-PCR | 22510686 |
| **hsa-miR-484** | SMAD2 | MIRT042141 | GFP reporter assay, Western blot | 28286418 |
| **hsa-miR-484** | ZEB1 | MIRT734111 | GFP reporter assay, Western blot | 28286418 |
| **hsa-miR-518d-3p** | PPA1 | MIRT438337 | Luciferase reporter assay | 24639097 |
| **hsa-miR-599** | MYC | MIRT733881 | Luciferase reporter assay, qRT-PCR, Western blot | 27398141 |
| **hsa-miR-599** | SATB2 | MIRT735048 | Luciferase reporter assay, Western blot | 28167280 |

**Supplementary Table 3:** Enriched KEGG pathways among miRNA target genes related to T2D processes and pathways

| **Name** | **Score** | **Expected Score** | **P-Value** | **Info** |
| --- | --- | --- | --- | --- |
| Insulin resistance | 3 | 0.245194 | 0.002246 | FOXO1, PIK3R3, SLC2A1 |
| Insulin signaling pathway | 5 | 0.311034 | 2.96E-05 | CRK, FOXO1, MAPK1, PIK3R3, RAF1 |
| Type II diabetes mellitus | 3 | 0.104435 | 0.000237 | CACNA1C, MAPK1, PIK3R3 |
| PI3K-Akt signaling pathway | 7 | 0.801423 | 2.86E-05 | BDNF, CDKN1A, EGFR, MAPK1, MYC, PIK3R3, RAF1 |
| Glucagon signaling pathway | 3 | 0.240654 | 0.002154 | FOXO1, SIRT1, SLC2A1 |
| AMPK signaling pathway | 4 | 0.272438 | 0.000237 | CCNA2, FOXO1, PIK3R3, SIRT1 |
